# Unsupervized Quantification of Demographic Structure for Single-copy Alignments

**DOI:** 10.1101/338442

**Authors:** AB Rohrlach, Nigel Bean, Gary Glonek, Barbara Holland, Ray Tobler, Jonathan Tuke, Alan Cooper

**Affiliations:** School of Mathematical Sciences, University of Adelaide, Adelaide, South Australia, 5005, Australia.; ARC Centre of Excellence for Mathematical and Statistical Frontiers, University of Adelaide, Adelaide, South Australia, 5005, Australia.; School of Natural Sciences (Mathematics), University of Tasmania, Hobart, Tasmania 7001, Australia.; Australian Centre for Ancient DNA, School of Biological Sciences, University of Adelaide, Adelaide, South Australia, 5005, Australia.

## Abstract

Single-copy sequence alignments have been a valuable source of information for genetic studies; their lack of recombination makes phylogenetic analyses tractable [1]. Specifically, mitochondrial DNA will continue to play an important role in genetic studies due to its high mutation rate and high copy per cell count of the molecule [2]. In this paper we develop a new method for the analysis of single-copy sequence data that simultaneously considers the relationships between sequenced individuals and positions of interest in the genome. We then show that tests for relationships between genetic information and qualitative and quantitative characteristics can be calculated. We motivate the use of our method with examples from empirical data.

## 1 Introduction

An important feature of any genetic analysis can be detecting whether a sample comes from a structured population. Demographic structure can take many forms. For example, samples may be taken from geographically isolated subpopulations [3], from subpopulations along a migration route [4] or from temporally separated population replacement events [5]. In some cases it can be of interest to discover that no geographic structure exists at all, leading to the exploration of social structure [6].

A popular form of unsupervized data exploration is principal components analysis (PCA) [7]. PCA is a dimension reduction technique that takes *n p*-dimensional vectors and, using linear combinations of the original vectors, finds *min*(*n* − 1, *p*) *p*-dimensional basis vectors. The new vectors are ordered by the amount of variability explained by each ‘principal dimension’. Often the first few dimensions are used to visualise points in the new transformed space.

PCA is a non-parametric, hypothesis-free exploratory technique, making it a particularly attractive analytical tool. However, PCA does require that the vectors of information are quantitative variables. Clearly sequence characters are not quantitative random variables, and so a transformation must be applied to raw sequence data before PCA can be directly applied [8]. However, we are aware of no such suitable transformation for DNA sequences that are non-biallelic, and in particular, haploid DNA such as mitochondrial DNA (mtDNA) or Y chromosome sequences. Instead, we suggest the application of multiple correspondence analysis (MCA) directly to the sequence characters.

MCA is an adaptation of PCA where categorical variables (in this case Single Nucleotide Polymorphisms: SNPs) are converted into binary variables denoting the presence or absence of each level of the variables (in this case alleles) [9]. Unlike PCA, MCA can be applied to any number of alleles. Our method makes the assumption that SNP inheritance is random, i.e. that the underlying phylogenetic tree is a star tree. One could test whether alleles appear to occur independently by investigating a contingency table of pairwise allele counts for an alignment, and then apply a chi-squared test. Since one would almost always overwhelmingly reject the null hypothesis, the result of a chi-squared test would be of no interest. However, the matrix of signed residuals under this assumption form the basis of the transformation from sequence data to continuous data.

In this paper we aim to show that MCA is a statistically powerful method for the analysis of non-autosomal DNA. We show that MCA has many properties that are analogous to PCA, and is hence immediately intuitive to researchers with experience using PCA. We demonstrate that PCA only quantifies the relationships between rows (individuals), while MCA quantifies the relationships between the rows (individuals) and also the columns (SNPs) simultaneously. For this reason we can quantify and visualise relationships between individuals as in PCA, and also quantify and visualise the relationship between SNPs, and between SNPs and individuals simultaneously in the same dimensions.

We show that results obtained from MCA correspond to the results obtained from mtDNA phylogenetic trees in a meaningful way, and that demographic structure can be detected using these results. We explore an alignment of African mtDNA from haplogroups L0, L1, L2, L4 and L5 to show that our method produces valid and easily interpretable results by reproducing mtDNA macro-haplogroups via clustering. We also explore an alignment of modern and ancient thylacine mtDNA from a mainland and island population. We show that thylacine genetic signals are highly correlated with longitude, and identify a possible ancestral migration route. Finally we explore an alignment of Western Australian Ghost Bat mtDNA to show that genetic diversity can be almost completely explained by discrete cave locations.

## 2 New Approaches

MCA is a generalised form of correspondence analysis that can be seen as the counterpart of PCA for categorical data analysis. Utilizing this powerful unsupervised data exploration method for genetic data yields a number of useful results and techniques that PCA does not allow.

First, the method may be applied directly to alignment data, forgoing the need for a transformation of sequence data to normalised allele frequency counts which inherently assumes data comes from a population at Hardy-Weinberg equilibrium [8]. Second, the method is able to calculate coordinates for individuals in genetic space (as in PCA) but can also simultaneously calculate coordinates for genetic markers (such as SNPs). Here we derive a multi-dimensional coordinate space scaling that allows the coordinates of individuals and SNPs to be directly visualized, and for demographic structure to be explored in both spaces simultaneously. Finally we define methods for exploring supplementary data in the case of both continuous and discrete variables, and show that the results of MCA can be used to identify SNPs of interest, leading to the detection of diagnostic SNPs, or potentially selective markers.

## 3 Results

### Coordinates in Gene Space and Dissimilarity Matrices for Haplotype Identification

The L-haplogroups represent the earliest evolution in modern human history, with the most recent common ancestor (MRCA) of the L-haplogroups being the MRCA of all humans. Hence, our method should be able to recover structure in the form of clusters of the major haplogroups L0, L1, L2, L4 and L5. To test this, we analysed a custom alignment from several published studies involving African sampled mtDNA [10, 11, 12, 13, 14]. We randomly chose sequences from these studies from sub-haplogroups L0d, L0k, L1c, L2a, L4 and L5a. We aimed to include 20 samples per haplogroup, although we included only 9 from L5a, as this was all that was available at the time of writing, and 10 from L4 to deliberately introduce further sampling asymmetry, resulting in an alignment of 79 individuals (see Table S1 for the file list of Genbank accession numbers and haplotype assignments).

We aligned our sequences to the revised Cambridge Reference Sequence [15] using **`MAFFT v7.310`** [16]. Haplogroups were determined using **`Haplogrep v2.1.0`** [17]. Aligned sequences were filtered to remove any homogeneous sites. MCA was performed on the remaining 281 SNPs. The first two principal dimensions captured 50.93% of the total inertia. That is, 50.93% of the variability in the 16,569 dimensional space (the number of base pairs in the sequences) can be observed in the first two principal dimensions.

We reconstructed a phylogenetic tree to compare the topology with our results. A Tamura-Nei model, with invariant sites and a gamma distribution with five classes was selected as the best model of sequence evolution using **`ModelGenerator v0.85`** [18]. We used **`Beast v1.8.3`** [19] to construct the phylogenetic tree using an MCMC chain of length 5x 10^9^, logging parameters every 10,000 states. The first 5 × 10^8^ states were discarded as burn-in, and the remaining trees were used to find a consensus tree using **`treeannotator v1.8.4`** [19]. Convergence was assessed through trace plots of posterior distributions. The branches of the consensus tree are in evolutionary time (relative mutation rate *μ* = 1) as we are only interested in the topology of the tree as a means of comparison with the results of the MCA (see Figure 1).

**Figure 1:**
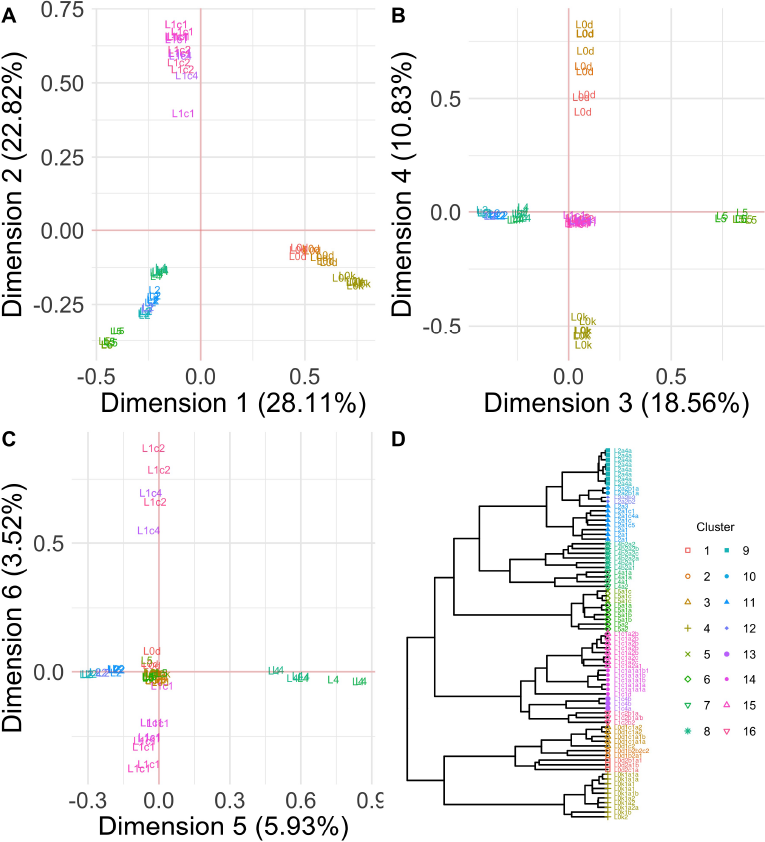
Scatter plots of the first six principal dimensions and phylogenetic reconstruction for the L-haplogroup alignment. Colors indicate cluster assignment via hierarchical agglomerative clustering.

In the first two principal dimensions (Figure 1, panel A), L0 (bottom right quadrant) is visibly separated from the remaining haplogroups, and this makes sense, with L0 being the most divergent human mtDNA haplogroup. The deep split within L0, between L0d and L0k can be observed here, with the 10 furthest points representing the L0k sub-haplogroup.

L1 then separates (top left quadrant) from L2, L4 and L5 (bottom left quadrant), which is the next major split in the human mtDNA tree. L5 is also separated from L2 and L4, and this is the next major split. Finally, although it is not as pronounced as the previous separations, L2 and L4 separate, and this is the final major split.

The third dimension (Figure 1, panel B) shows a clear distinction between L5 (positive coordinates) and L2 and L4 (negative coordinates). The fourth dimension (Figure 1, panel B) separates L0d (positive) coordinates from L0k (negative coordinates). Dimension 5 (Figure 1, panel C) finds a separation between L2 (negative coordinates) and L4 (positive coordinates). Finally dimension 6 (Figure 1, panel C) separates L1c1 from the remaining L1c individuals. The remaining dimensions further identify splits in the tree, though this is not included in Figure 1. For this reason, when performing clustering we include all principal dimensions.

We performed hierarchical agglomerative clustering on the coordinates from the MCA using the R-package **`cluster v2.0.6`** [20]. The choice of termination point for identifying clusters is arbitrary, and in our case we cease identifying clusters when a cluster of size one is suggested.

The clustering algorithm respected the configuration of the points in the first two principal dimensions. The first cluster identified was L0, followed by L1 and L5. L0d and L0k are separated into two clusters, followed by the split between L2 and L4, which reflects the greater divergence time for the respective haplogroups [21]. The fifth cluster separation of L2 and L4 represents the final major haplogroup according to the current nomenclature.

The remaining clusters all respect the sub-haplogroup structure of the mtDNA tree, identifying sub-haplogroups for each of L0, L1, L2, L4 and L5. The clusters identified here are specific to this dataset, *i.e.* it may be the case that if more than one sequence from the haplogroup L1c2b2 were included, then we may have identified L1c2b2 as a cluster.

It is worth noting that our clustering suggests that the current nomenclature for human mtDNA may not reflect statistically significant groups, but rather just the sequence of historically discovered diagnostic SNPs in densely sampled haplogroups. For example, the split between L0d and L0k appears more significant than the split between L2 and L4 in both the MCA and the phylogenetic tree. However, the sample sizes here are not large enough to refute the nomenclature, although the method provides a clear way forward to revise this.

Overall, the method has clearly shown that we can identify a tree like structure in the data, and that just the first two principal dimensions were able to visualise the haplotype structure in the data.

### Application of method for continuous supplementary variables

The thylacine (*Thylacinus cynocephalus*) is an Australian marsupial carnivore most famous for its recent extinction due to human hunting [22, 23]. By the time of the arrival of Europeans to Australia, the thylacine had already undergone a significant population decline, was extinct on the mainland and was only found in Tasmania.

From museum samples we use sequence data from three samples from south-west Western Australia (WA), three samples from the Nullarbor Plain in WA, six samples from Tasmania (TAS) and one sample from New South Wales (NSW) (see Figure 2) [23]. Samples were removed if the longitude or latitude were unknown, or if the sampling age was unknown. To avoid artificial inflation of signal from geographical coordinates, for sequences found in the same location, a single representative was randomly selected. In total, 13 individuals were analysed (see Table S2 for supplementary variables and Genbank accession numbers).

**Figure 2:**
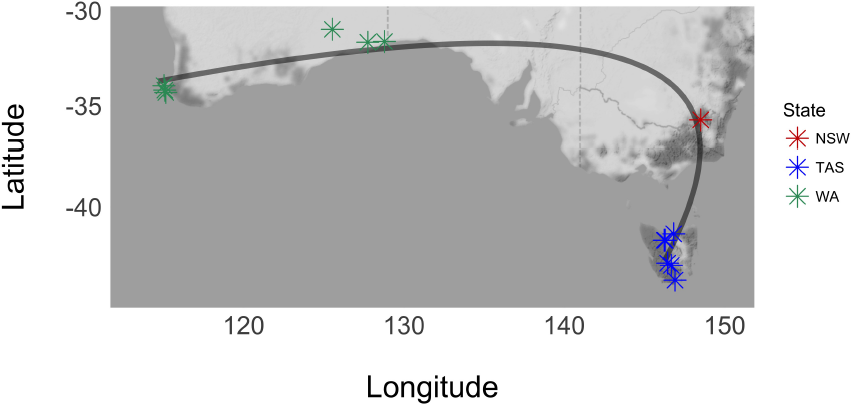
Sample location and sample IDs for thylacine mtDNA. The black line is the predicted geographic locations for thylacines given the observed range of principal dimension 1 coordinates. Colours indicate the location in which samples were found (Red=NSW, Blue=TAS, Green=WA).

Sequences were aligned using MAFFT v7.310 [16]. The alignment was filtered to remove homogeneous and missing sites, and a total of 113 SNPs were included in the MCA.

From the MCA row factor scores the first principal dimension, which captured 62.62% of the total inertia, correlates strongly with longitude (*r* = 0. 9467235, *p* = 9.517 × 10^−7^), suggesting a possible migration gradient [24]. Gradients are not expected to be strictly linear for principal component maps, and the same can be assumed for MCA maps [4].

To investigate the relationship between geography and the MCA coordinates, a multi-response linear model was used. Multi-response linear models are similar to standard linear models, but allow for more than one response variable to be collectively modelled by the same set of explanatory variables [25]. A multi-response model was fitted to the data to predict latitude and longitude using principal dimension 1 (PD1). Polynomial models of varying degrees were fit and the best model was quadratic (using AIC), with *R*^2^ values of 0.9334 and 0.9075 between longitude and latitude respectively.

A ‘predicted’ migration route can be projected onto the geographical map suggesting a coastal route was taken along the south of Australia (see Figure 2). However, the extremely small sample size means the results are limited as the MRCA of the sample is not necessarily closely related to the MRCA of the population, and there is little reason to believe that ancestral thylacine populations remained in the areas they originally inhabited.

### Application of method for categorical supplementary variables

The ghost bat (*Macroderma gigas*) is a native Australian bat endemic to the Northern Pilbara and Kimberley in Western Australia, and in some regions of the Northern Territory and Queensland [26]. The ghost bat conservation status is currently listed as vulnerable by the International Union for Conservation of Nature.

Ghost bats are found in discrete populations within cave-based colonies. Blast mining disrupts, and in some cases destroys, cave complexes, displacing resident bat populations. Conservationists wish to understand the phylogeographic distribution of ghost bats to understand if the destruction of a single cave colony significantly reduces genetic diversity. Gene flow between colonies would indicate a reduced impact on ghost bat diversity, whereas a highly structured population would indicate a need to reduce the effects of blast mining and colony disruption.

We focus on samples collected in the Northern Pilbara found in four colonies on the northern side of the Hamersley Range (Bamboo, Callawa, Lalla Rookh, Klondyke), and two on the southern side (Rhodes, Silvergrass), and one colony in the Kimberley (Tunnel Creek) (see Figure 3). The Hamersley Range contains the twenty highest peaks in Western Australia, and so forms a significant geographical boundary for bats to cross. All colonies are represented by one sampled cave, with the Rhodes colony being the exception with four closely sampled caves (see Table S3 for supplementary variables and ID numbers).

**Figure 3:**
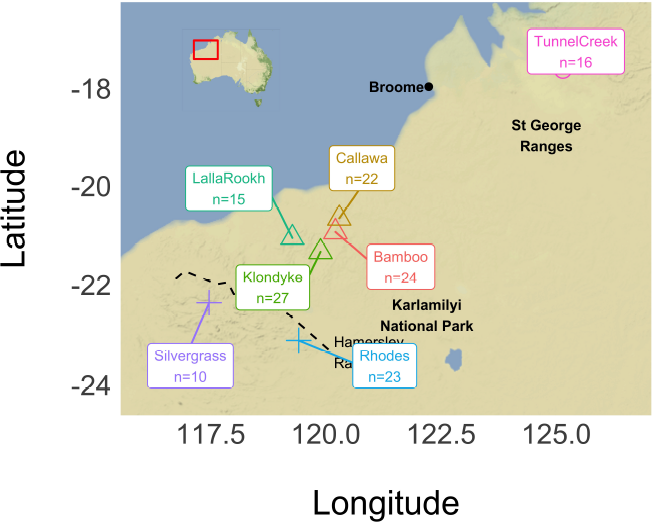
Sample location and sizes for ghost bat mtDNA. Colors indicate colonies and and shapes indicate

We filtered an alignment of 257bp of the mtDNA HVR region, from 137 individuals, to remove homogeneous sites, and MCA was performed on the remaining 25 SNPs. For each individual, the colony and population (North, South, Kimberley) were recorded and treated as categorical supplementary variables. Longitude and latitude were also recorded and kept as quantitative supplementary variables.

In Figure 4 we present the squared-correlation plot for all supplementary variables and SNPs. The *x* and *y* coordinates of points in this plot give squared correlation values for the first two principal dimensions and each of the supplementary variables and SNPs. The further to the right of the plot a variable or SNP name is, the more highly correlated it is with Dimension 1. Similarly, the further to the top of the plot a variable or SNP is, the more highly correlated it is with Dimension 2. Immediately we see that latitude and longitude are not as strongly correlated with the two first principal dimensions as population and colony.

**Figure 4:**
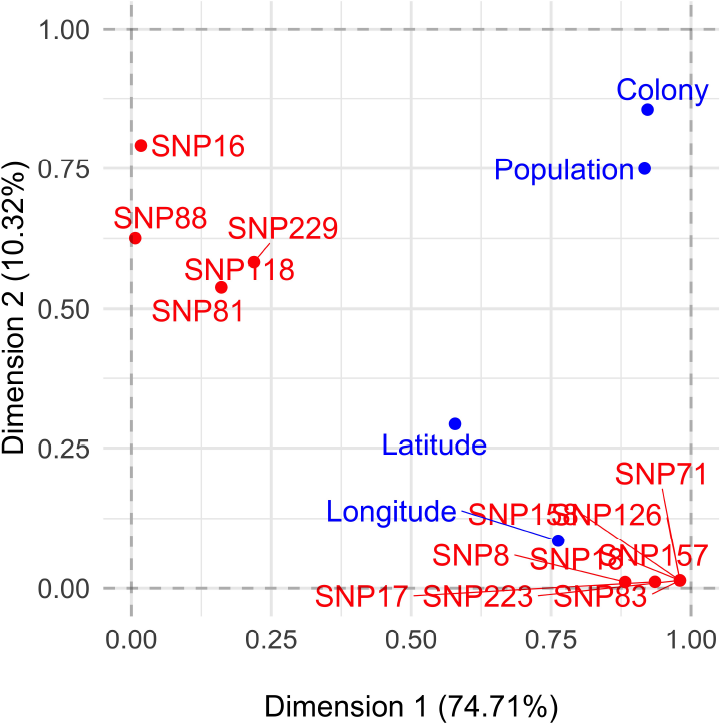
The correlation plot for all variables and SNPs for the Ghost Bat alignment.

Calculating the *η*^2^ values for population structure and colony structure yields 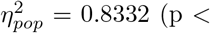 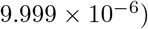 and 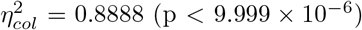 respectively.

Clearly then, population explains a large proportion of the variability of the points in genetic space, however colony explains a greater proportion of the total variance, and thus had a larger *η*^2^ value. Clearly colony explains a significant proportion of the structure of the individuals in genetic space since the first two principal dimensions explained a total of 85.03% of the total inertia.

The first principal dimension visualises the split between the Kimberley and Pilbara colonies (except for one Pilbara individual within the Kimberley samples), and the second principal dimension visualises the divide between the North and South colonies within the Pilbara region. In fact, if one places a boundary representing the Hamersley Range, on the y-axis at −0.35 (the dashed line in Figure 5), only one individual from the Northern Pilbara sample lies below the boundary, and only one Southern Pilbara sample lies above the boundary.

Since 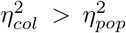, this suggests that colony better explains the structure of the genetic coordinates.

**Figure 5:**
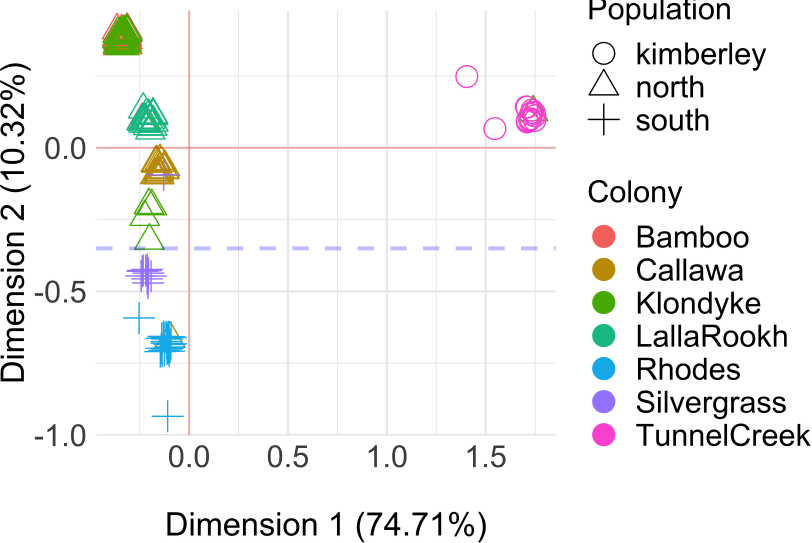
The scatter plot of the first two principal dimensions for the ghost bat alignment.

This can be seen in Figure 5 where we observe that five of the seven colonies form distinct clusters, with the exception of: a single Klondyke individual in the Tunnel Creek colony, a Callawa individual in the Rhodes colony, and a Silvergrass individual in the Callawa colony. The remaining two colonies, Klondyke and Bamboo, cluster together in the top left of the plot. A further four individuals from Klondyke form a separate cluster, genetically nearer the southern colonies. This may represent a recent migration from the southern colonies, or potentially members of the founding population for the southern colonies.

Without further information we cannot explain why these two colonies cluster together. It is worth noting that the Bamboo Creek site is a recently abandoned complex of mines (dismantled in 1962), whereas mining operations in the Klondyke mine area have increased drastically since 1955. It is possible that there has been a recent blending of the two colonies as bats have been displaced from places like Klondyke, and found refuge in caves left from abandoned mining efforts, like Bamboo Creek. However, our results still indicate a significant colony-based structure outside of these two colonies. This colony-based structure further strengthens the argument for more protection for roosting colonies from blast mining.

Finally we give an example of identifying potential diagnostic SNPs using our method. From Figure 4 we also see that we can identify potentially diagnostic SNPs. Given that Dimension 1 is explained by the geographical separation of the Kimberley and Pilbara populations, as all but one of the individuals with a positive first principal dimension coordinate are from the Kimberley, SNPs that are highly correlated with this dimension may be diagnostic for one of the regions. For example, SNP18 can be found in only Kimberley ghost bats (with the exception of the one Klondyke individual). If we had decided to use clustering to identify haplogroups, then SNP18 could be considered a diagnostic SNP, from this limited sample.

## 4 Materials and Methods

### The transformation of genomic data to continuous coordinates

Consider an *n* × *p* alignment *A* of mtDNA, where *A*_*ij*_ ϵ {*A*,*C*,*G*,*T*}, filtered to remove homozygous sites. The *n* rows represent sequenced individuals, denoted {*a*_1_, ⃜, *a*_*n*_} and the *p* columns represent single nucleotide polymorphisms (SNPs), denoted {*s*_1_, ···, *s*_*p*_}. Note that each of the SNPs can take between two to four forms, and we say that *s*_*j*_ has |*s*_*j*_| levels.

Consider each of the 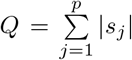 different allelic forms of the *p* SNPs, ordered (without loss of generality) numerically by position, then within SNPs, lexicographically by nucleotide. We can define an *n* × *Q* indicator matrix *X*, such that *X*_*ik*_ equals one if individual *a*_*i*_ has the allele at the position indicated by the *k*th column name, for *k* = 1, ···, Q (see Figure 6). Note that for a SNP with |*s*_*j*_ | levels, there are only |*s*_*j*_| − 1 linearly independent columns of information in the *X* matrix (since if an individual does not have any of the first |*s*_*j*_| − 1 forms of the allele, they must have the remaining allele). Hence, in total there are only *Q* − *p* linearly independent columns. Finally, we can also calculate a contingency table of pairwise marker combinations *B* = *X*^*T*^*X* (see Figure 6), this matrix is discussed later in the process.

**Figure 6:**
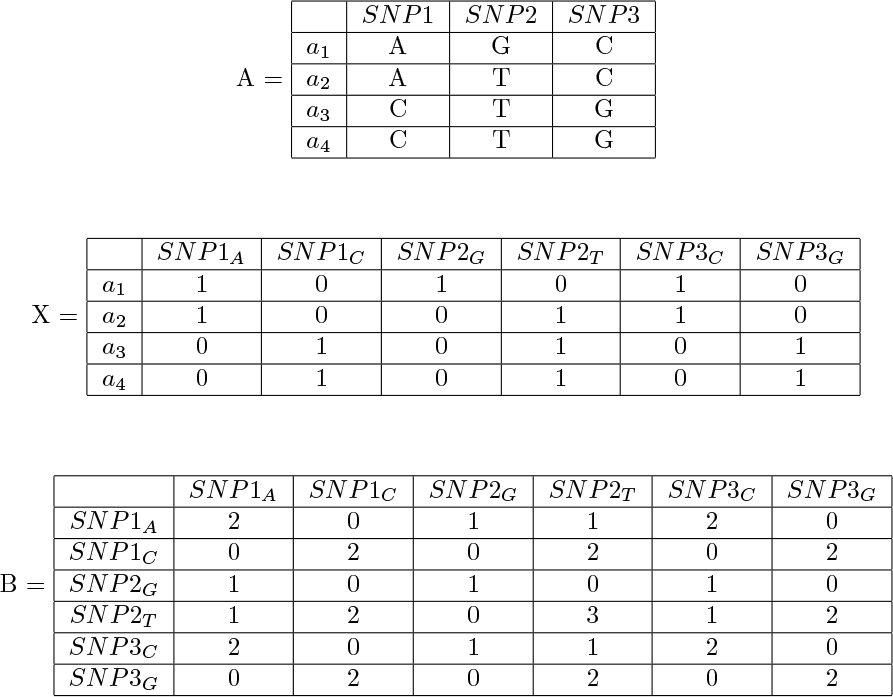
A transformation from raw sequence alignment *A*, to an indicator matrix *X*, and a Burt table *B* = *X*^*T*^*X*.

Let 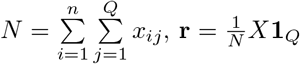, 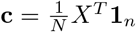 and 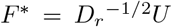 where 1_*k*_ is a *k* × 1 vector of ones, and define *D*_*r*_ = diag(*r*) and *D*_*c*_ = diag(*c*). We can define a new *n* × *Q* matrix, as a function of *X*, 
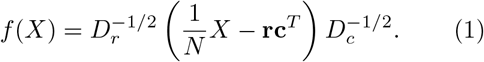

On *f* (*X*) we perform a compact singular value decomposition (SVD) so that *f*(*X*) = *U*σ*V*^*T*^. Due to the above number of linearly independent columns, the diagonal matrix of singular values, σ, will only have *J* = *Q* − *p* non-zero entries, and we need only consider these dimensions. Following this reasoning, *U* and *V* are truncated to be matrices of dimensions *n* × *J* and *Q* × *J*, respectively. From the diagonal matrix σ we may also obtain the percentage of inertia (analogous to variability in PCA) explained by each of the first *J* principal dimensions, which are proportional to the singular values.

The *standard* row and column coordinates, defined as *F*\* = *D*_*r*_^−1/2^*U* and 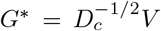 respectively, are the unscaled row and factor scores that do not account for the proportion of inertia in principal dimensions. A natural choice for scaling the standard row coordinates is to post multiply each dimension by the associated singular value. Hence the relative spread of points in each dimension is proportional to the amount of inertia captured by each dimension.

From the standard row scores we obtain the transformed coordinates, also called the ‘row factor scores’, and denoted *F*, of the individuals in the alignment *A* in ‘genetic space’ via 
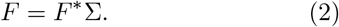

The distances between individuals calculated form these coordinates will respect three properties:

1. If two individuals have the same DNA sequence, they will have identical coordinates.
2. If two individuals share many alleles, they will be closer than two individuals that do not.
3. Individuals that share rare alleles will be closer still.

It is important to note that the pairwise distances between individuals calculated from the matrix *F* differ from classical pairwise genetic differences in two important ways. First, one need not assume a model of sequence evolution to find the matrix *F*. Second, classical pairwise genetic distances are calculated on only two sequences at a time, and so do not take into account the rarity of alleles. Our method uses the complete alignment to calculate the matrix *F*, and gives greater weight to rarer alleles.

The choice of rescaling for the standard column coordinates, with respect to the standard row coordinates, depends on the desired properties of the resulting column factor scores. We propose rescaling the standard column coordinates by the squares of the singular values, such that the column factors scores are 
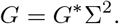

This rescaling of the standard column coordinates yields a desirable property for comparing the coordinates of individuals and alleles. The coordinates for any allele can be found at the centroid of the coordinates of the individuals that carry that allele (proof given in Appendix A). A special case of this property is that if an individual uniquely carries an allele, then that allele shares exactly the same coordinates as the individual (proof given in Appendix A).

There is a second, equivalent way to consider the method we have proposed. It is known that the row and column factor scores from *B* = *X*^*T*^*X* will be the same as the factor scores obtained from *X* [27]. The transformation *f* (*B*) (found in a similar same way as in Equation 1, but with appropriate dimensions for r, c and a recalculated normalising constant) would yield a *Q* × *Q* matrix, of the form *R* = [*ρ*_*ij*_]. *R* is a matrix of the correlations for detecting linkage disequilibrium with multiple alleles, where if *ρ*_*ij*_ ≠ 0, then the loci associated with alleles *i* and *j* are in linkage disequilibrium [28]. While mtDNA does not undergo recombination, our method also attempts to identify groups of alleles that occur together more than expected just by random chance, and hence the individuals that carry these alleles.

As with PCA, we can use the principal coordinates to visualize the relationships between individuals. However, our method also allows us to visualize the relationships between SNPs, and between individuals and SNPs. We can also look at the pairwise distances between individuals, and SNPs, in genetic space.

Figure 7 shows the relationship between the sequences as shown in Figure 6. Since *a*_3_ and *a*_4_ have identical sequences, they have the same coordinates in gene space. As ai shares no similarity with *a*_3_ or *a*_4_, they are the furthest apart. However, *a*_2_ shares one SNP with *a*_3_ and *a*_4_, and two SNPs with a single individual *a*_*i*_, and hence is more closely ‘attracted’ to *a*_1_. Due to this ‘attraction’ to individuals with similar SNP profiles, the term ‘inertia’ is used in the place of ‘variance’.

Note the relationship between individual coordinates, and SNP coordinates. Since *a*_3_ and *a*_4_ are the only individuals with ‘SNP1_C’ and ‘SNP3_G’, they share the same coordinates (the same can be said of *a_1_* and ‘SNP2_G’). ‘SNP1_A’ is shared by *a*_1_ and *a*_2_, and so falls exactly at the mid-point of the two points. However, ‘SNP2_T’ is shared by *a*_2_ and by both *a*_3_ and *a*_4_, and so lies only one-third the way along the line connecting *a_3_* and *a*_4_ to *a*_2_.

**Figure 7:**
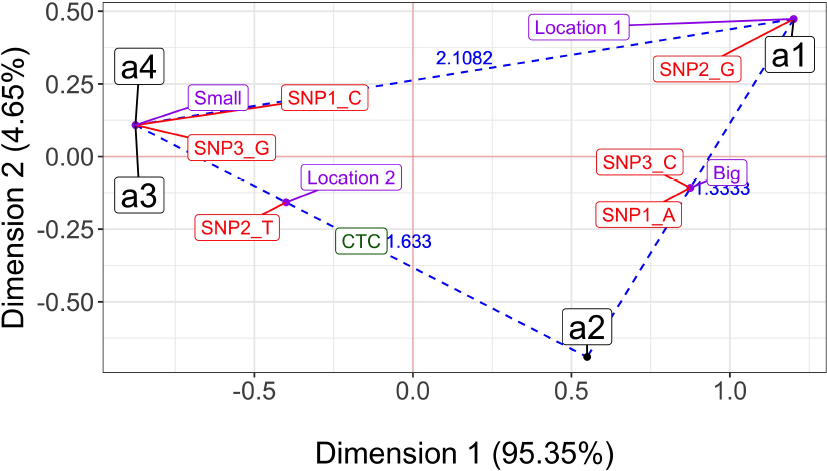
A biplot of the first two principal dimensions for the sequences as shown in Figure 6. Individuals are in black, SNPs are in red and projected coordinates for supplementary variable Size (Big and Small) and Location (Location 1 and Location 2) are shown in purple. The new sequence ‘CTC’ is projected onto the dimensions and given in green. Euclidean distances between individuals are given in blue.

Note that in Figure 6, if an individual has an ‘A’ at the first site, then they always have a ‘C’ in the third position. Similarly, if an individual has a ‘C’ at the first site, then they always have a ‘G’ in the third position. Hence the SNPs at the first and third sites provide no new information about the nature of the relationships between individuals since one can infer the third SNP, given the nature of the first SNP. For this reason, the first two principal dimensions capture 100% of the inertia, and reducing the dimensionality of the transformed genetic space results in no loss of information about the structure of the relationships between individuals.

It is possible to project new sequences onto the genetic space defined by an MCA. The new sequence must have one of the allelic forms for every SNP from the original alignment. For example, consider an alignment of new sequences of dimension *m* × *p* denoted *H*, with corresponding *m* × *Q* indicator matrix *i*_*H*_ (see Figure 8).

For the matrix *s*_*H*_ = diag (*i*_*H*_ 1_*Q*_), coordinates for the new sequences can be found [29] via 
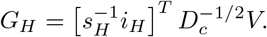

In Figure 7 we project the new sequence given in Figure 8 onto the first two principal dimensions as given by the analysis of the alignment A from Figure 6. Note that this sequence is an equal ‘mix’ of the sequences *a*_2_ and *a*_3_, and so falls halfway along the line connecting the two sequences. While this makes mathematical and intuitive sense, this mixing of two sequences makes no sense for non-recombining DNA.

**Figure 8:**
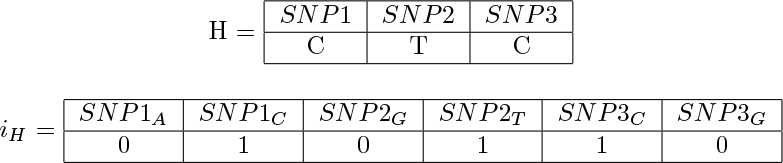
A transformation from a new raw sequence alignment *H*, to an indicator matrix *i*_*H*_ to be projected onto existing MCA dimensions.

While this method could be used for pseudo-haploid DNA where this type of interpretation does make sense, there is value in projecting new sequences onto the principal dimensions for non-recombining DNA. For example, when data contains many SNPs, principal dimensions may represent haplogroups with a collection of diagnostic SNPs. Projecting ancient samples, for example, would include individuals ancestral to individuals from the alignment that have not acquired more recent SNPs. These projected points might fall along the line connecting the origin to the group, with individuals that carry fewer diagnostic SNPs closer to the origin.

Finally, we may project the ‘average’ coordinates of some qualitative supplementary variable. Imagine we have *r* such variables, with a total of R levels. Let W be the *n* × *r* matrix of supplementary information, with corresponding *n* × *R* indicator matrix *j*_*W*_ (see Figure 9).

Following a similar method for projecting new sequences, for the matrix *s*_*W*_ = diag(1_*RjW*_), coordinates for the average qualitative supplementary variables can be found via 
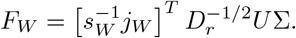

The projected coordinates for the supplementary variables are an estimate of the average coordinates for individuals with the given levels of the qualitative supplementary variables. For example, in Figure 9, if you were to imagine that we could sample all individuals from the true population with level ‘Big’ for variable ‘Size’, then we believe that the centre of the cluster of points would fall at approximately (0.875, −0.1083). Note that the calculated coordinates are based on inertia, and called ‘barycentres’ rather than centroids.

**Figure 9:**
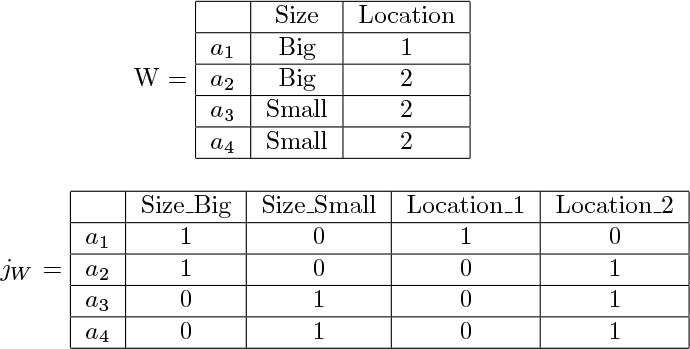
A transformation from a matrix of supplementary qualitative information *W*, to an indicator matrix *jw* to be projected onto existing MCA dimensions.

In Figure 7 we see that only sequences *a*_3_ and *a*_4_ have Size ‘Small’, and since they share coordinates, the barycentre for ‘Small’ also shares this coordinate. Sequences ai and *a*_2_ both have Size ‘Big’, and so the barycentre for ‘Big’ falls halfway along the line connecting their coordinates. Similarly, *a*_1_ is the only individual found at ‘Location 1’, and so the barycentre for ‘Location 1’ shares the coordinates of *a*_1_. However, *a*_2_, *a*_3_ and *a*_4_ were all found at ‘Location 2’, and so the barycentre for ‘Location 2’ can be found two thirds of the way along the line connecting *a*_2_ to *a*_3_ and *a*_4_. Notice also that ‘Location 2’ is found exclusively when there is a T at SNP2, and so they also share coordinates.

Once a coordinate representation of the relationship between individuals has been constructed, we can examine relationships between individuals in this ‘genetic space’, and compare them to characteristics that have been recorded for individuals. These ‘supplementary variables’ are anything recorded about sampled individuals that were not used in the alignment table (*i.e.* any non-SNP data). Of particular interest are demographic variables, such as country of origin, spatial coordinates on a landscape or morphological characters, for example.

Here we give three examples that illustrate the ability of the method to produce biologically meaningful and intuitive results in a rigorous statistical framework, using previously published empirical data sets.

### Correlation tests for continuous supplementary variables

Identifying relationships between coordinates in genetic space and continuous supplementary variables is intuitively simple. One could simply calculate the Pearson correlation coefficient for each continuous supplementary variable, followed by an exact test for a significantly non-zero coefficient.

It should be noted for a principal components analysis of spatially structured sequence data that has undergone recombination, that the top two principal components are expected to be highly correlated with perpendicular geographic axes [4]. In the case of mtDNA, or any other recombination-free sequence data, this assumption cannot be made. More extreme axis values can be interpreted as the accumulation of more and more of a unique set of SNPs that characterize some partition of the most-related tips of a tree.

### Correlation tests for categorical supplementary variables

Supplementary categorical variables which explain significant proportions of the structure of individuals in gene space can be identified, and their effect quantified, using the correlation ratio *η*^2^ [30]. The correlation ratio *η*^2^ can be thought of as the proportion of variability explained by a qualitative variable. For a one-dimensional response variable *Y*, *η*^2^ is equivalent to *R*^2^, the coefficient of determination for a linear model with the qualitative variable as the sole predictor variable. In the case of multiple dimensional response variables, an analogous *η*^2^ may be calculated [31].

A permutation test can be used to find if *η*^2^ is significantly greater than for a random relabelling of the population. For each of the *T* permutations of the group labellings, we calculate 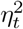, the correlation ratio calculated for the *t*^*th*^ permutation. An empirical p-value of the form (*r*+ 1)/(*T* +1) is calculated, where *r* is the total number of permuted samples yielding a greater correlation ratio than the observed sample [32].

## Discussion

MCA provides a powerful method for unsupervized exploration of single-copy DNA. Our method is analogous to a PCA analysis of classical allele correlation values for detecting linkage disequilibrium. We have shown that *p*-dimensional single-copy DNA can be transformed into coordinates in genetic space, analogous to the way in which diploid DNA is transformed via PCA in many genetic studies. One of the attractive features of our approach is the parallel with PCA, making the interpretation of results natural for researchers experienced with PCA.

Our method allows for the coordinates of supplementary variables to be calculated and visualized in the same coordinate space as for individuals and SNPs, and for the relationships between the supplementary variables and principal dimensions to be quantified. Like PCA, additional sequences can be projected onto the coordinate space that has been calculated from an alignment of interest.

Dimension reduction can be performed, reducing potentially massive numbers of SNPs into far fewer dimensions with potentially little reduction in information, leading to informative visualization of highdimensional data. Unlike PCA, our method is able to simultaneously investigate the relationships between individuals and SNPs. This extra information can lead to the detection of diagnostic SNPs, and potentially SNPs that are correlated with supplementary variables of interest such as habitat or phenotypic traits.

Our method was able to detect known haplotype structure, showing that the results of MCA are biologically meaningful. Similarly, the fact that our method can be reformulated as a PCA of the linkage disequilibrium table for multiple loci indicates that applications to recombining DNA may also be useful for detecting population structure.

Using techniques from classical statistics, our method was also able to efficiently visualise the strength of the relationships between supplementary information and empirical sequence data. Finally, using standard polynomial regression techniques, our method was able to identify a possible migration route for geographically distributed sequence data.

## 4.1 Appendix A

We aim to show that our choice of scaling for the row and column factor scores yields the property that if an individual uniquely carries an allele, then the individual and the allele share the same coordinates.

To do this we investigate properties of the indicator matrix *X*, where the columns have been permuted to make the first column the identifying allele and to make the first row the identified individual (without loss of generality). To avoid carrying constants, we assume that *X* has already been normalized to have grand sum one.

### Result 1

If there is an allele that uniquely identifies an individual, then the individual and the allele have the same coordinates if the standard row factor scores are scaled by the singular values, and the standard column factor scores are scaled by the squared singular values.

**Proof:** Let *X* be an *n* × *Q* matrix such that 
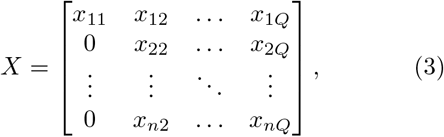
such that 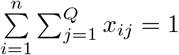 and *x*_*ij*_ ≥ 0 ∀*i, j*.

Let 1_*k*_ be a *k* × 1 vector of ones, and let 
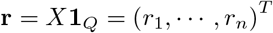
and 
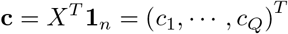
be the strictly positive row and column sums of *X* respectively, and let 
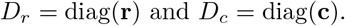

We begin by showing that the SVD of the matrix 
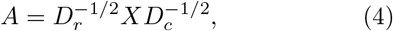
has a particular structure. We then exploit this to show the required result for the matrix 
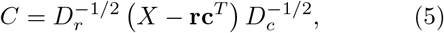
namely that for our choice of row and column standard coordinate scaling, the identifying allele and the identified individual share the same scaled factor scores.

Let *A* have a compact singular value decomposition (SVD) of the form 
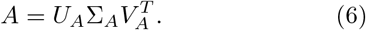

Consider the matrix product 
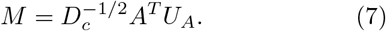

Since 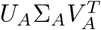 is a SVD, then *U*_*A*_ is a unitary matrix, and so 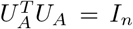 (where *I*_*k*_ is the *k* × *k* identity matrix), and substituting Equation (6) into Equation (7) gives, 
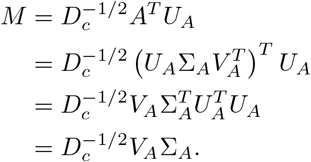

Hence, the first row of *M* is 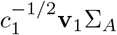, where v_1_ is the first row of *V*_*A*_.

However, instead substituting Equation (4) into Equation (7) yields 
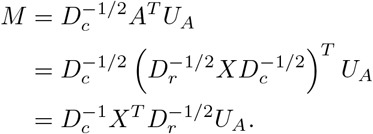

Note that since *x*_21_ = ··· = *x_n1_* = 0, then *c*_1_ = *x*_11_ and hence the first row of 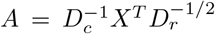 will equal 
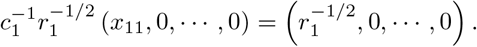

So the first row of *M* is also 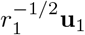, where u_1_ is the first row of *U*_*A*_.

This shows that 
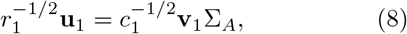
which are the first rows of the row and column scores of the SVD of 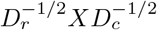, where *X* is of the form given in Equation (3).

To extend this result to the SVD of the matrix *C* (SVD) of the form in Equation 5, first note that 
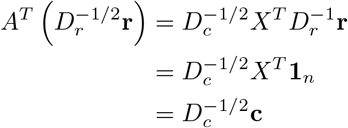
and 
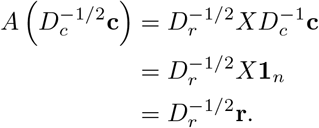

Now this shows that the SVD of *A* has a singular value of 1, with left and right singular vectors 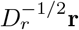 and 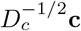, respectively.

We now show that a SVD of *C*, denoted 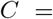
 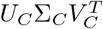 can be augmented by these singular vectors and the singular value 1 to construct a SVD for *A*.

Consider new matrices 
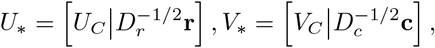
and 
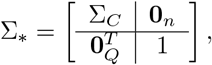
where 0_*k*_ is a *k* π 1 vector of zeros. Next we show that *U*\*, *V*\* are unitary matrices and π* is a rectangular matrix diagonal matrix with non-negative real numbers on the diagonal. 
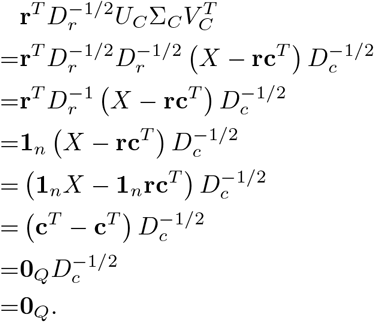

Since, σ_*C*_ is a diagonal matrix with positive diagonal entries, we know that 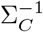 exists. Further, 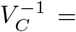 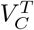, so this implies that 
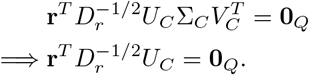

Similarly, it can be shown that 
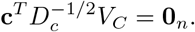

It follows then that 
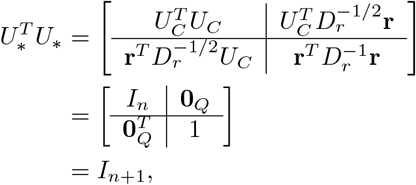
and that 
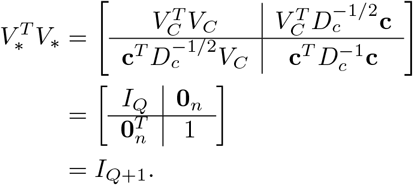

Note that 
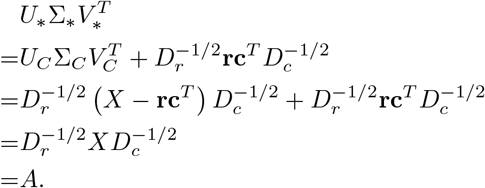

Therefore, since π_*_ is a rectangular diagonal matrix, with positive diagonal entries, and *U*_*_ and *V*_*_ are unitary matrices, it must be that 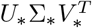 SVD of *A*.

Thus we have two representations for the compact SVD of *A*. Hence they are equivalent. However, from Equation 8, we know that the first row and column factor scores for the SVD of *A* are equal, and are given by 
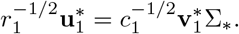

Hence any sub vectors that are constructed from removing corresponding elements from the vectors 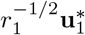 and 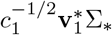 will also be equal, specifically 
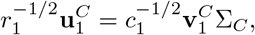
where 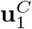 is the first column of *U*_*C*_, and 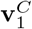 is the first column of *V*_*C*_.

### Result 2

If a single allele identifies a group of *m* individuals, the the column factor score for the allele is the centroid of the row factor scores of the identified individuals, if the standard row factor and the standard column factor scores are scaled by the squared singular values.

**Proof:** If 
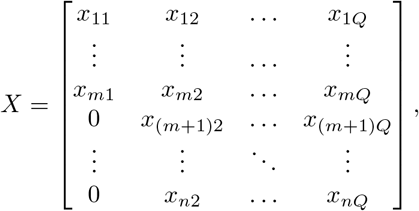
where *x*_11_ = *x*_*m*1_ > 0, and *x*_(*m*+1)1_ = *ηηη* > *x*(*n*1) = 0, and 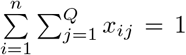 and *x*_*ij*_ ≥ 0 ∀*i, j*. As previously, the first column of 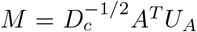 is 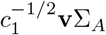. However, the sum of the first column of *X* is *mx*_11_, and hence the first row of 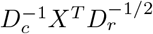 is now 
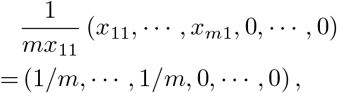
and it follows that the first column of *M* is also 
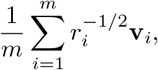
yielding that 
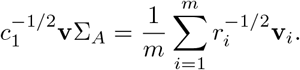

Following the same argument as before, it is also true that this is the case for the SVD of 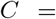 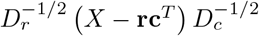. Hence the identifying allele has column factor score equal to the centroid of the row factor scores for the identified individuals.

